# CRISPR-mediated Correction of Oncogenic AS-NMD in Splicing Factor Mutant Cancer

**DOI:** 10.64898/2026.07.28.739648

**Authors:** Preeti Nagar, Md Rafikul Islam, Nirjhor Anuvob Rahman, Shabiha Afroj Heeamoni, Md Mahbub Hasan, Saaimatul Huq, Rezwan Ali, Mokarram Hossain, Mohammad Alinoor Rahman

## Abstract

Alternative splicing coupled to nonsense-mediated mRNA decay (AS-NMD) evolved as a master regulator of gene expression. Dysregulated AS-NMD has been identified as the root of many human maladies, from developmental defects to deadly cancer. Poison exons (PEs) are highly conserved alternative exons that contain a premature termination codon and elicit AS-NMD when included in a transcript. Cancer cells often exploit the inclusion of PEs to downregulate tumor suppressors or the exclusion of PEs to upregulate oncoproteins. Therefore, PEs have drawn significant attention as a novel therapeutic avenue for cancer and other diseases. Here, we examine a therapeutic proof-of-concept for manipulating PE-mediated oncogenic AS-NMD using a CRISPR-based approach. Using paired guide RNA, we successfully deleted a PE of a tumor suppressor (EZH2) from the genome of SRSF2-mutated leukemia. This editing resulted in EZH2 mRNAs without a PE, escaped AS-NMD, and restored the protein expression. This subsequently reinstated H3K27 histone methylation and rescued defective chromatin regulation associated with impaired hematopoietic stem cell differentiation. Finally, we showed the preferential advantages of CRISPR over the antisense technology we recently developed targeting the PE of EZH2. Therefore, the CRISPR strategy shows compelling evidence as a therapeutic approach targeting PE in cancer and other human diseases.

## Introduction

The human genome evolved as a highly coordinated blueprint that preserves precise instructions for the appropriate regulation of gene expression, enabling diverse biological functions and responses to internal and external challenges. Alternative splicing (AS) and nonsense-mediated mRNA decay (NMD) are two post-transcriptional RNA processing mechanisms. They play crucial roles in regulating gene expression (1–3). AS promotes proteome diversity using a limited number of genes. NMD evolved as a surveillance mechanism to eliminate faulty transcripts with a premature translation termination codon (PTC), such as those generated by nonsense or frameshift mutations. AS also often generates mRNAs with a PTC, which are degraded by NMD. This coordinated RNA processing mechanism is called AS coupled to NMD (AS-NMD). Studies in recent decades have highlighted AS-NMD as a powerful regulator of gene expression, playing a pivotal role during development, cell proliferation and differentiation, metabolism, immune surveillance, and many other biological processes (4, 5).

About 95% of human multiexon genes undergo AS, and ∼33% of alternatively spliced transcripts are targets for NMD (6–8). Dysregulated AS-NMD has been identified as the root of many human maladies, including developmental defects, neurological diseases, metabolic disorders, immune dysfunction, genetic abnormalities, and deadly cancer (4, 5, 8–11). AS-NMD can be exploited positively or negatively to drive pathological consequences. For example, AS-NMD can compromise cellular integrity by degrading proteins essential for physiological functions, immune surveillance, and tumor suppression. In contrast, evading or suppressing AS-NMD can compromise gene expression fidelity and may allow the expression of defective or pathogenic proteins, such as oncoproteins. Poison exons (PEs) are a class of highly conserved cassette exons that comprise PTCs and therefore promote AS-NMD when included in transcripts (12–14). Although poison exons are not used for protein-coding in host genes, they play an important role in controlling gene expression (5, 14, 15). For example, the serine-arginine-rich (SR) splicing factors (SFs) contain ultra-conserved PEs in their genes (SRSF1-SRSF12) (12–14). Although SR proteins are important for normal splicing regulation, their altered expression can turn them into oncoproteins (16–19). In normal cells, when SR protein levels are upregulated, SR proteins autoregulate their expression through AS-NMD by including PEs (12–14), thereby avoiding an effect in oncogenic transformation.

Tumor cells often exploit PEs to differentially regulate AS-NMD in their favor. SR proteins are frequently overexpressed in various solid tumors (4, 16, 20–22), including breast, colon, lung, ovarian, liver, and pancreatic tumors. Recent studies have shown that upregulation of SR proteins in solid tumors is often correlated with reduced inclusion of PEs (14, 15). In contrast to splicing factor overexpression, recurrent mutations in splicing factors (most frequent in SRSF2, SF3B1, U2AF1, and ZRSR2) are frequently identified in hematologic malignancies, such as myelodysplastic syndromes and leukemia, and are also common to different solid tumors (4, 23, 24). It has been reported that several important genes linked to hematopoiesis are downregulated by the inclusion of PEs (4, 25–30). Therefore, PEs have received significant attention in recent years as potential therapeutic targets due to their strong links with various cancers and numerous human diseases.

Strategies to manipulate PE-mediated AS-NMD are emerging in genetic diseases and cancer (4, 5, 31, 32). However, therapeutic efficacy is often compromised by technological limitations. Antisense oligonucleotide (ASO) pharmacology is a popular strategy to modulate AS or AS-NMD (32, 33). However, the efficacy of ASO is often compromised by delivery challenges, suboptimal dosing, stability issues, or low throughput. RNA-mediated interference (such as siRNA or shRNA) can’t change alternative splicing in a gene-specific manner. Small molecules often appear toxic due to off-target effects and fail in clinical trials (31, 34, 35). These disparities remain as critical roadblocks to therapeutic success. Therefore, research should focus on developing novel, effective targeted therapeutic approaches grounded in a comprehensive understanding of molecular regulations.

Here, we examine a gene-specific targeted approach to modulate PE-mediated oncogenic AS-NMD using CRISPR/Cas9. Although CRISPR/Cas9 is frequently used to knock out a target gene, our strategy restores its expression by preventing the inclusion of a PE, thereby evading AS-NMD. As a proof-of-concept, we investigated a blood cancer model harboring an oncogenic mutation in the splicing factor SRSF2. We and others previously showed that mutations in SRSF2 (SRSF2^Mut^) promote aberrant inclusion of a PE in the tumor suppressor gene *EZH2* and elicit AS-NMD (25–27). This results in downregulated EZH2 protein levels, altered chromatin regulation, defective hematopoiesis, and leads to the development of cancer (25). To challenge these molecular maladies, we exploited the CRISPR/Cas9 system with paired guide RNA (pgRNA) targeting the flanking intronic sequences of the PE of *EZH2* in SRSF2^Mut^ model cancer cell lines. Genomic deletion of the poison exon resulted in correction of the splicing error of *EZH2,* evading AS-NMD, and restoring the protein expression. This subsequently enhanced H3K27 histone methylation and rescued defective chromatin regulation. This proof-of-concept shows promising therapeutic potential targeting PE-mediated AS-NMD in cancer and many other human diseases.

## Materials and methods

### Cell culture

SET2, MOLM-13, TF1, K052, K562, and K562P95H cells were cultured in RPMI-1640 medium with 1% penicillin streptomycin (Sigma) and supplemented with 10% fetal bovine serum (FBS) and maintained in an incubator at 37°C with 5% CO_2_. HEK 293T cells were cultured in DMEM medium with 1% penicillin streptomycin (Sigma) and supplemented with 10% fetal bovine serum (FBS) and maintained in an incubator at 37°C with 5% CO_2_.

### Splicing analysis using reverse transcription PCR (RT-PCR)

Total RNA was isolated using TRIzol reagent, followed by DNase treatment. Reverse transcription reaction was conducted using ImProm-II reverse transcriptase (Promega). During cDNA synthesis, an oligo (dT) reverse primer was used. PCR reaction was performed by Phusion High-Fidelity PCR Master Mix with HF Buffer (M0531L, New England Biolabs). The primers for *EZH2* were: 5’-TTTCATGCAACACCCAACACT-3’ and 5’-CCCTGCTTCCCTATCACTGT-3’. PCR products were separated on a 2% agarose gel and stained with SYBR Safe (Invitrogen). The gel image was captured using the ChemiDoc MP Imaging System (Bio-Rad). PCR bands were quantified, and the percent spliced-in (PSI) ratio was calculated as the exon-included isoform band intensity divided by the intensity of included and skipped isoform bands.

### Protein expression analysis by western blotting

For protein isolation, cells were washed and harvested in 1x PBS containing Protease Inhibitor Cocktail. After centrifugation at 2,000 x g for 5 min, the cell pellets were resuspended in buffer A [10 mM HEPES-NaOH (pH 7.8), 10 mM KCl, 0.1 mM EDTA, 1 mM DTT, 0.5 mM PMSF, 0.1% Nonidet P-40, Protease Inhibitor Cocktail], followed by incubation on ice for 30 min. The samples were then sonicated and centrifuged at 20,000 x g for 5 min. The supernatants were collected as total cell lysate for western blotting. The antibodies used were: EZH2 (Cell Signaling, 5246), GAPDH (Cell Signaling, 2118), Histone H3 (Cell Signaling, 4499), and Tri-Methyl-Histone H3 (Lys 27) (Cell Signaling, 9733).

### Designing of pgRNA and cloning

pgRNAs located in or near the targeted exon were designed according to the concept described previously (36), using both NAG and NGG protospacer-adjacent motifs. Candidate pgRNAs were cloned into the Lenti-multi-CRISPR vector (Addgene#85402) following a published method (37). All plasmids were sequenced using Sanger sequencing to confirm the desired sequences.

### Lentiviral transduction

HEK 293T cells were cultured in DMEM medium with 10% FBS with antibiotics. The next day, the medium was replaced with fresh DMEM containing 10% FBS, chloroquine diphosphate, and no antibiotics. Cells were transfected with the Lenti-multi-CRISPR-pgRNA vector, pMD2.G, and psPAX2, and incubated at 37°C for 6 hours. The medium was replaced with fresh DMEM supplemented with 10% FBS and without antibiotics. After 48 hours, the lentiviral supernatant was collected and filtered through 0.45 mm filters. K052 or K562^Mut^ cells were grown in RPMI-1640 medium with 10% FBS. The next day, K052 or K562^Mut^ cells were transduced with lentiviral supernatant containing polybrene, and the procedure was repeated with fresh virus the following day. The next day, cells were cultured in fresh medium supplemented with 1 μg/mL puromycin for 8 days, with the medium replaced every 2 days. Cells were then split into 96-well plates for single-clone expansion. Positive single clones were selected and confirmed following PCR and Sanger DNA sequencing of the genomic DNA.

### EZH2 inhibitor

EPZ-6438 (Tazemetostat) was selected as an EZH2 inhibitor (EZH2i), purchased from SelleckChem (Cat# S7128), and used according to the manufacturer’s instructions.

### Antisense oligonucleotides (ASO)-mediated splicing correction

We synthesized ASO from Integrated DNA Technologies (IDT). The ASO2 sequences: 5’-TCCAACAGGCAATAT-3’ (38). We transfected K052 cells with ASO2 using Lipofectamine 3000 (Invitrogen). RNA or proteins were isolated 48 h later after the transfection.

### Quantification and statistical analysis

Densitometric quantification of RT-PCR or western blotting was performed using ImageJ. Statistical analysis was performed either by t-test or ANOVA. Data are presented as mean and standard deviation and indicated by error bars. Graphs were generated using GraphPad Prism software.

## Results

### AS-NMD is a crucial mediator of tumorigenesis in splicing-factor-mutated cancers

Heterozygous mutations in splicing factors have been recurrently identified in cancer patients with myelodysplastic syndromes (MDS), acute myeloid leukemia (AML), chronic lymphocytic leukemia (CLL), and a variety of solid tumors (23). The most frequent mutations occur in SF3B1, SRSF2, U2AF1, or ZRSR2. These mutations typically alter RNA-binding properties, leading to genome-wide splicing alterations (25–30, 39–46). Interestingly, certain mRNAs promoted by different mutant splicing factors comprise PTCs and are degraded by AS-NMD. Some of these events are linked with tumorigenesis and are critical vulnerable targets for cancer therapy (4, 19, 21, 47).

We previously analyzed the Cancer Genome Atlas AML patient RNA-sequencing data set (TCGA-LAML) with or without SRSF2 mutation (SRSF2^Mut^ vs. SRSF2^WT^) (26). We reanalyzed the data to include a potential oncogenic AS-NMD target for our investigation. Among 843 differential splicing events (DSE), the majority of the events were alternative cassette exon splicing (CE), and the others included alternative splice site selection (ASS) and alternative intron retention (IR). Among these DSE, ∼16% were AS-NMD targets. Again, the majority of the AS-NMD events represent CE (**Fig. 1A-B**). Experimental validation of tumorigenic activity identified only a handful of targets, considered as cancer driver events, while others are passenger splicing events with no functional relevance to cancer development (25–27, 48). Several consistent, noteworthy, and validated cancer driver events include: a poison-exon inclusion-mediated AS-NMD in *EZH2;* an intron-retention-mediated AS-NMD in *INTS3;* and an alternative-exon-exclusion-mediated AS-NMD in *CLK3* (25–27, 48). EZH2 (Enhancer of Zeste 2 Polycomb Repressing Complex 2 Subunit) is an enzyme that catalyzes methylation of histone H3 at lysine 27 (H3K27) and plays an important role in chromatin regulation (49), suppression of gene expression, stem cell differentiation and development, and determination of cell fate (49) (**Fig. 1C**). Downregulation of EZH2 protein expression through AS-NMD, promoted by SRSF2^Mut^, drives defective hematopoietic differentiation progressing to cancer (25–27). INTS3 is a member of the integrator complex 3, which functions in snRNA processing and RNA polymerase pause release. Downregulation of INTS3 protein expression through AS-NMD, driven by SRSF2^Mut^, promotes increased stalling of RNA polymerase II (RNAPII) (27). CLK3 (CDC-like kinase 3) is a protein kinase that regulates pre-mRNA splicing by phosphorylating SR splicing factors. Aberrant skipping of an exon in *CLK3* in SRSF2^Mut^ cancer generates splicing-derived neoantigens (48). Since the focus of our proof-of-concept is modulating the splicing of oncogenic poison/cassette exon using CRISPR/Cas9, we chose *EZH2* in SRSF2^Mut^ blood cancer as our model oncogenic target.

**Figure 1.**
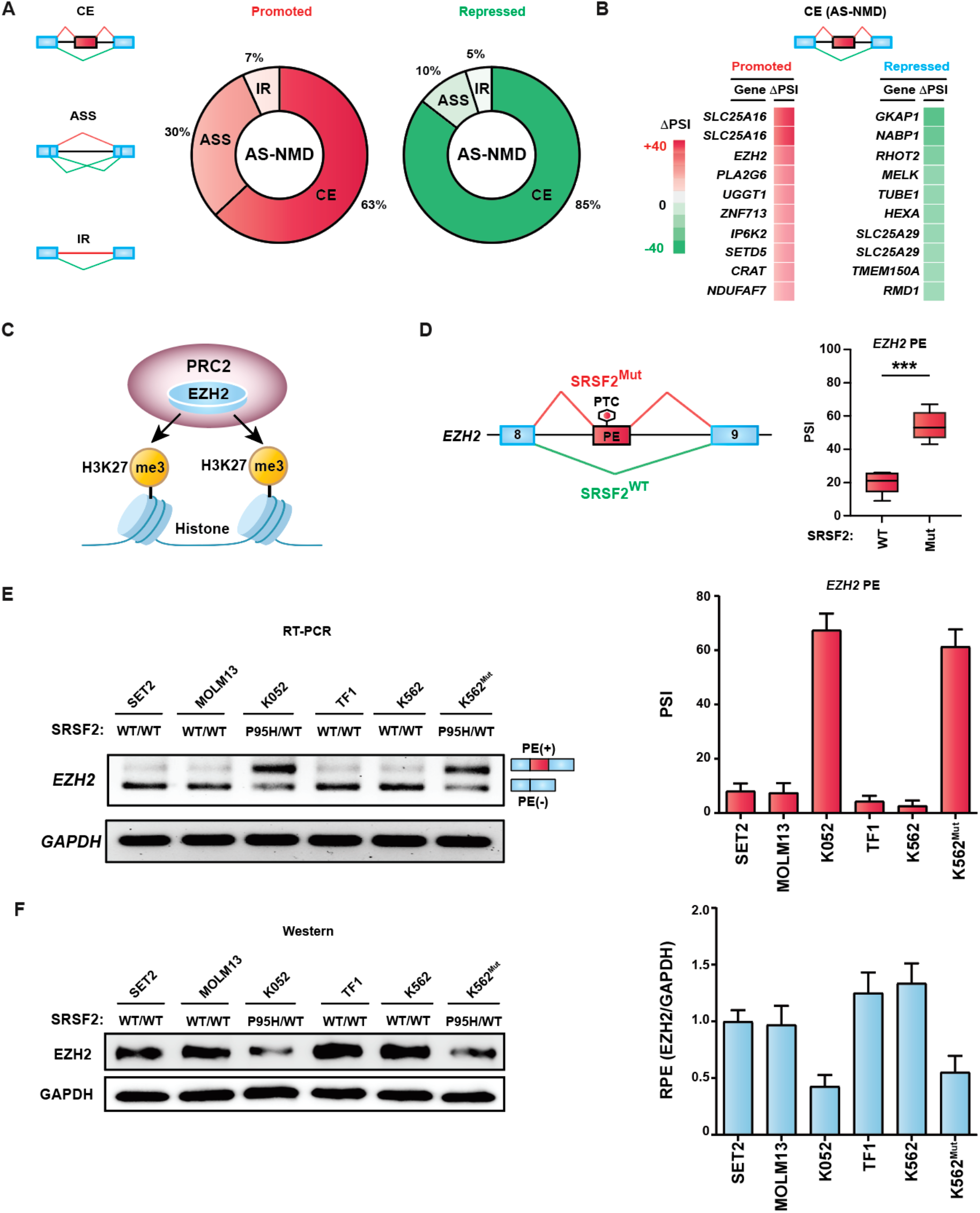
Genome-wide altered splicing profile promoted by oncogenic Pro95 mutations in SRSF2. **(A)** Proportion of differential AS-NMD events (promoted and repressed) analyzed from TCGA-LAML RNA-seq of AML patients (SRSF2^Mut^ vs. SRSF2^WT^) (26). ASS: alternative splice site selection; CE: cassette exon splicing; IR: intron retention. **(B)** Ranking of the top ten AS-NMD targets in promoted and repressed CE events, respectively. 1′PSI: differences in percent spliced-in (SRSF2^Mut^ vs. SRSF2^WT^). **(C)** Cartoon showing the function of EZH2 in histone methylation and chromatin regulation. PRC2: polycomb repressing complex. **(D)** Left, diagram showing *EZH2* poison exon (PE) splicing. Right, quantification of the *EZH2* PE inclusion as percent spliced-in (PSI) in TCGA-LAML samples from the genotypes indicated shown at the right (mean ± SD, n=5, ***p<0.001, t test). **(E)** Left, representative RT-PCR gel showing the *EZH2* PE splicing in different human leukemia cell lines from the indicated genotypes. Right, quantification of the *EZH2* poison exon inclusion as PSI. Bar graph (mean ± SD, n=3). **(F)** Left, representative western blotting showing indicated protein expression in different human leukemia cell lines from the indicated genotypes. Right, quantification of the relative protein expression (RPE) of EZH2 normalized against GAPDH (EZH2/GAPDH). Bar graph (mean ± SD, n=3).

### Recapitulation of aberrant inclusion of the *EZH2* PE in SRSF2^Mut^ cancer models

TCGA-LAML data showed exclusive inclusion of the *EZH2* PE in AML patients with SRSF2^Mut^, compared with SRSF2^WT^ (**Fig. 1D**). We also previously confirmed this in RNA samples from AML patients (38). This was also consistently identified in several other studies (25–27, 50, 51). To recapitulate this aberrant splicing in human cancer cell lines, we examined several leukemia-derived cell lines. We tested SET2, MOLM-13, K052, TF1, and K562 cell lines. Among these cell lines, SET2, MOLM-13, TF1, and K562 have wild-type SRSF2 genotype (SRSF2^WT/WT^), whereas K052 harbors a naturally occurring heterozygous P95H mutation in SRSF2 (SRSF2^P95H/WT^). RT-PCR showed exclusive inclusion of the *EZH2* PE in K052, but not in the other cell lines (**Fig. 1E**). In contrast, western blotting showed downregulation of EZH2 protein expression in K052 compared to other cell lines (**Fig. 1F**). We included an isogenic K562 cell line carrying a heterozygous SRSF2 knock-in mutation (SRSF2^P95H/WT^), termed K562^Mut^. We found increased inclusion of the *EZH2* PE in mRNA and concomitant downregulation of EZH2 protein levels in K562^Mut^ compared to K562 (**Fig. 1E-F**). These results clearly demonstrated that the PE-mediated AS-NMD of *EZH2* was promoted by SRSF2^Mut^. Therefore, K052 and K562^Mut^ were included as reliable model mutant cell lines for downstream studies.

### Testing the feasibility of CRISPR/Cas9 strategy to manipulate a cassette exon splicing in SRSF2^Mut^ cancer cells

To manipulate the expression of an oncogenic mRNA isoform toward the canonical and functional isoform by selectively deleting an oncogenic cassette exon, we exploited a previously described CRISPR/Cas9 approach termed paired guide RNA (pgRNA) for alternative exon removal (36, 52). In this approach, two guide RNAs targeting distinct regions flanking the target exon or a splice site of the target exon are delivered simultaneously into cells to enforce genomic deletion of the target region (**Fig. 2**). To test the feasibility of this approach in SRSF2^Mut^ blood cancer cells, we first sought a tractable exon that could be followed up with a chemical assay. We selected the *HRPT1* gene, encoding Hypoxanthine Phosphoribosyl Transferase 1 (36, 52). This enzyme is crucial for the purine salvage pathway, recycling guanine and hypoxanthine to form nucleotides. Inactivation of HRPT1 results in resistance to 6-thioguanine (6-TG) (36). We employed two sets of pgRNA targeting the constitutive exon 2 of *HPRT1*: pgHPRT1-S1 directing to the downstream splice site of exon 2; and pgHPRT-S2 directing the flanking intronic sites of exon 2 (**Fig. 3A**). We cloned these guide RNAs into the Lenti-Multi-CRISPR vector (Addgene) harboring Cas9 and puromycin resistance genes (**Fig. 2A**). We used the empty vector as a control. We then delivered the individual vectors into patient-derived K052 cells harboring a natural mutation in SRSF2 (SRSF2^P95H/WT^) through lentiviral transduction and selected positive clones after puromycin selection. We then plated positive clones for single-clone expansion. PCR of genomic DNA of the expanded clones showed the desired deletion in the majority of cases, with few exceptions (**Fig. 3B**). Sequencing of genomic DNA revealed that most clones have complete genomic DNA excision as expected (**Fig. 3C**), whereas others harbor diverse short insertions/deletions (indels). We used clones with complete genomic deletion for further analyses. Reverse transcription polymerase chain reaction (RT-PCR) of extracted RNA from the positive clones confirmed the skipping of *HPRT1* exon 2 in mRNA (**Fig. 3D**). Western blotting of extracted protein from the positive clones also confirmed inactivation of HPRT1 at protein levels (**Fig. 3E**). Furthermore, inactivation of HPRT1 through skipping of exon 2 resulted in the expected 6-TG resistance (**Fig. 3F**). These data suggest that pgRNA-mediated alternative exon removal can be exploited reliably in SRSF2^Mut^ blood cancer cells for mechanistic and biological studies.

**Figure 2.**
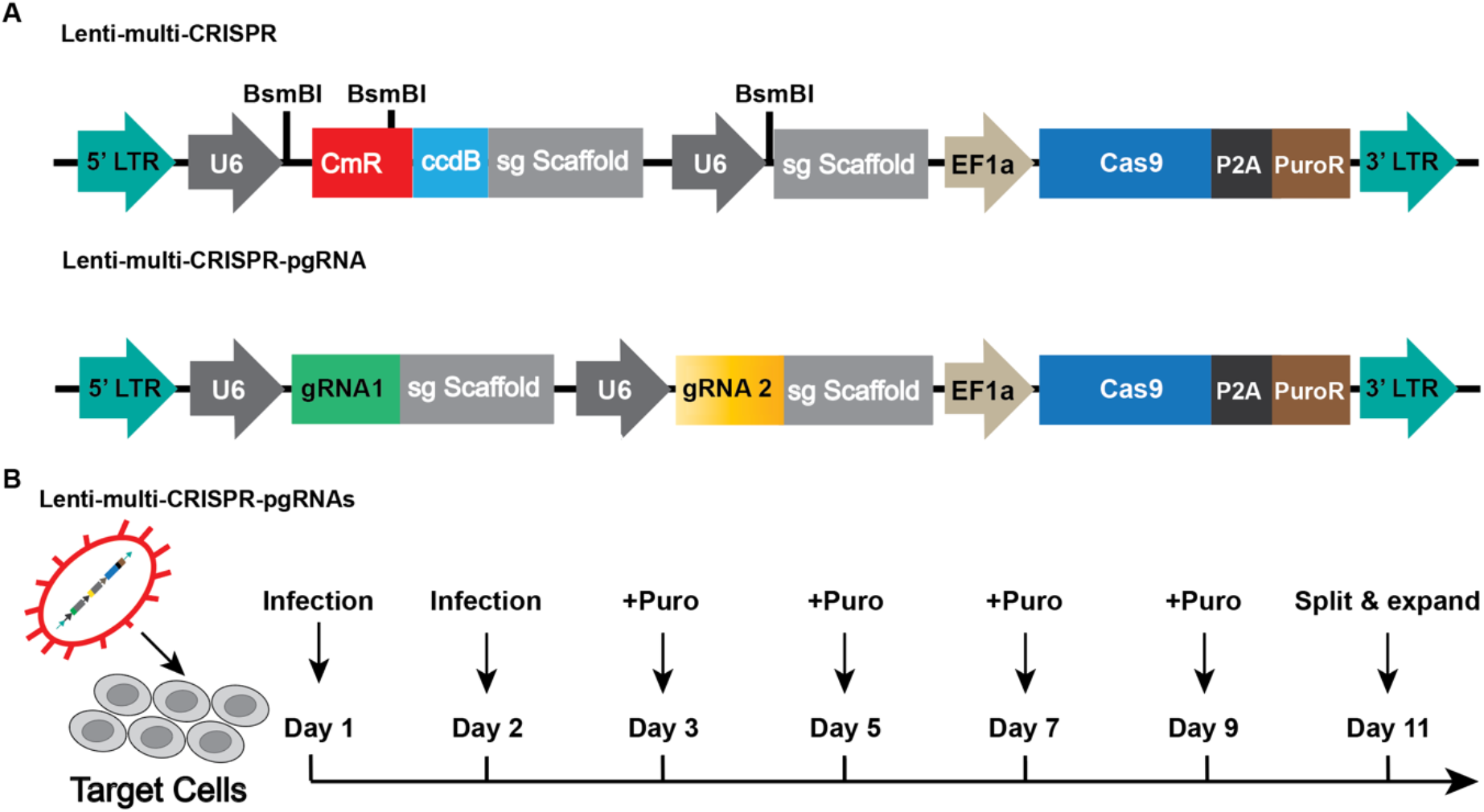
Strategy of pgRNA-mediated deletion of an alternative cassette exon. **(A)** Schematics of Lenti-multi-CRISPR and Lenti-multi-CRISPR-pgRNA vectors. LTR: long terminal repeat; U6: U6 promoter; CmR: Chloramphenicol resistance gene; ccdB: ccdB toxin gene; sg Scaffold: sgRNA scaffold sequence; EF1a: EF1 promoter; Cas9: Cas9 gene; PuroR: puromycin resistance gene. **(B)** Overview of the strategy of pgRNA delivery and puromycin selection.

**Figure 3.**
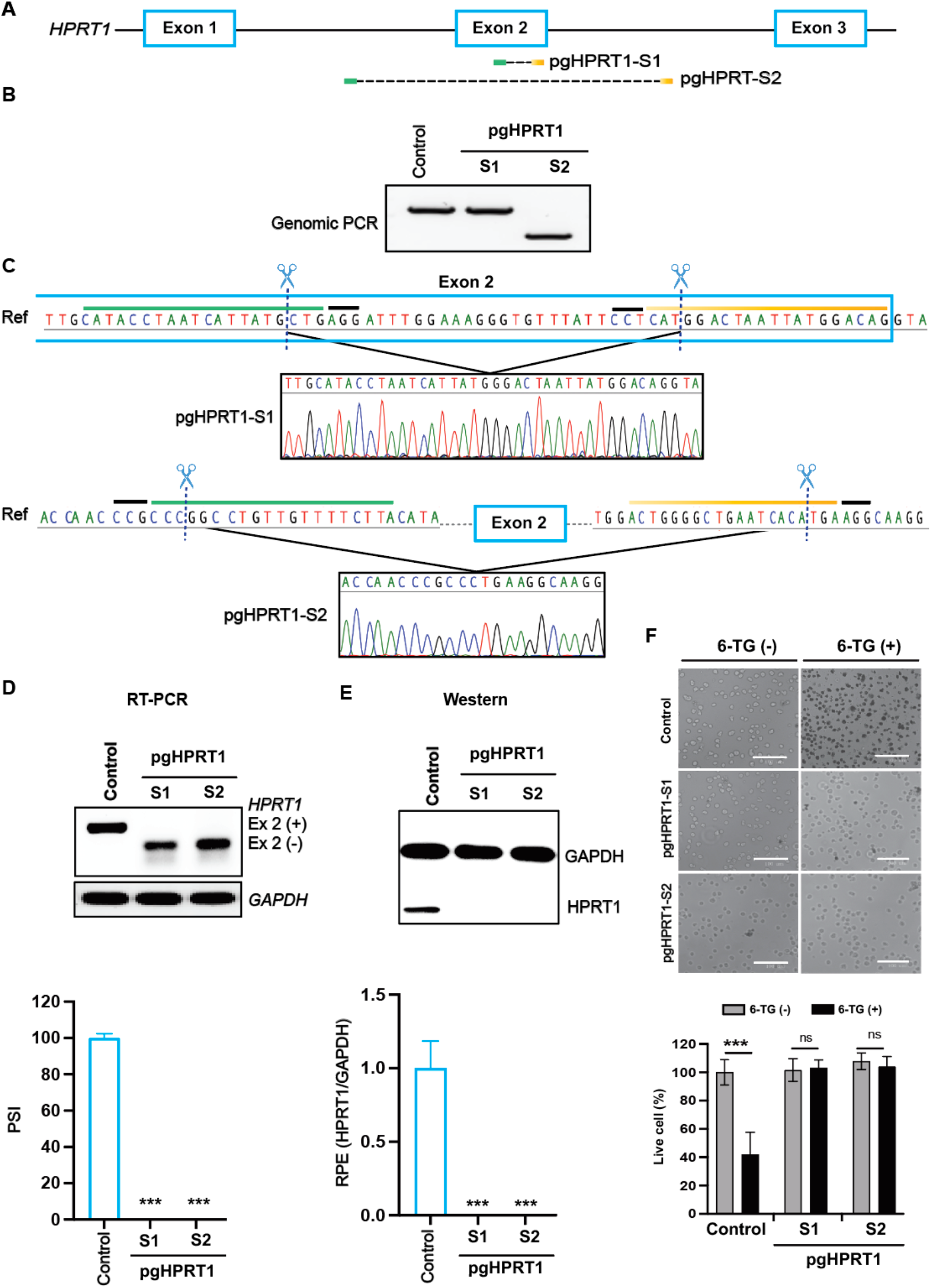
Evaluating the feasibility of the pgRNA-mediated exon deletion approach in an SRSF2-mutated model cancer cell line targeting a tractable exon of *HPRT1* using a chemical assay. **(A)** Schematics of two sets of pgRNA (S1 and S2) to delete genomic sequences targeting exon 2 of *HPRT1*. **(B)** Representative gel showing the genomic PCR of control (empty vector) or pgRNA-edited K052 clones targeting the *HPRT1* exon 2. **(C)** Representative Sanger sequencing of pgRNA-edited K052 clones showing sequence deletion targeting the *HPRT1* exon 2. **(D)** Top, representative RT-PCR gels of control (empty vector) or pgRNA-edited K052 clones showing the mRNA profiles of *HPRT1* and *GAPDH*. Ex2 (+): *HRPRT1* mRNA isoform with exon 2; Ex2 (-): *HRPRT1* mRNA isoform without exon 2. Bottom, quantification of the *HPRT1* exon 2 inclusion in mRNA as percent spliced-in (PSI). Bar graph (mean ± SD, n=3, ***p<0.001, t test). (**E**) Top: representative western blotting of control (empty vector) or pgRNA-edited K052 clones using antibodies against GAPDH and HPRT1. Bottom, quantification of the relative protein expression (RPE) of HPRT1 normalized against GAPDH (HPRT1/GAPDH). Bar graph (mean ± SD, n=3, ***p<0.001, t test). (E) Top, a phase-contrast image (scale: 100 µM) of control (empty vector) or pgRNA-edited K052 clones treated without (-) or with (+) 6-TG. Bottom, quantification of the live cells (%). Bar graph (mean ± SD, n=3, ***p<0.001, ns: not significant, t test).

### Genomic deletion of the *EZH2* PE from SRSF2^Mut^ cancer cells using pgRNA

To delete the *EZH2* PE from the genome of K052, we designed three sets of pgRNA (pgEZH2-S1, pgEZH2-S2, and pgEZH2-S3) directing the flanking intronic sites of the PE (**Fig. 4A**). We cloned the sequences for these pgRNA sets into the Lenti-Multi-CRISPR vector and generated Lenti-Multi-CRISPR-pgRNA vectors (**Fig. 2**). We used the empty vector as a control. We then transduced K052 (SRSF2^P95H/WT^) cells with these vectors. After puromycin selection, we processed positive clones for single-clone expansion. We didn’t observe any noticeable morphological changes in K052 cells after pgRNA treatment (**Fig. 4B**). PCR of genomic DNA of the expanded clones showed deletion of the expected size for most of the clones (**Fig. 4C**). Sequencing of genomic DNA revealed that most clones have complete genomic DNA excision as expected (**Fig. 4D**), whereas others harbor diverse short indels. We also tested the strategy in a second model cell line, K562^Mut^, with a knock-in mutation in SRSF2 (SRSF2^P95H/WT^), using the same three sets of Lenti-Multi-CRISPR-pgRNA vectors and following a similar approach. Again, we observed no noticeable morphological changes in K562^Mut^ cells after pgRNA treatment (**Fig. 4E**). PCR and sequencing of genomic DNA from the expanded clones also showed successful excision of the desired genomic segment in most cases, along with some short indels (**Fig. 4F-G**). We included clones with complete genomic deletion as desired for downstream analyses.

**Figure 4.**
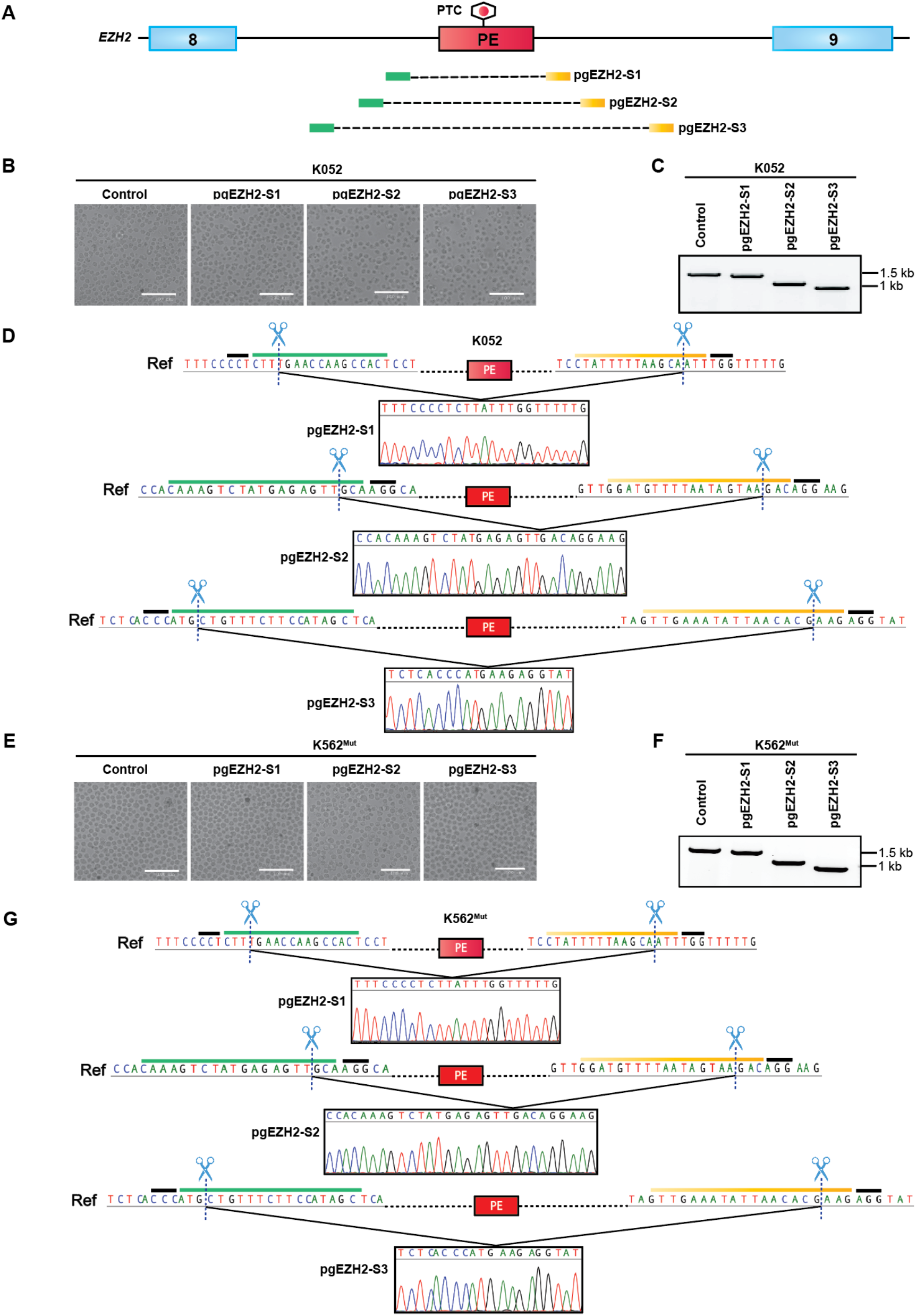
pgRNA-mediated genomic deletion of the *EZH2* PE in SRSF2-mutated model cancer cell lines. **(A)** Schematics of three sets of pgRNA (S1, S2, and S2) targeting the flanking intronic sequences of the *EZH2* PE. **(B)** Phase-contrast images of control (empty vector) or pgRNA-edited K052 clones. **(C)** Representative gel showing the genomic PCR of control (empty vector) or pgRNA-edited K052 clones targeting the *EZH2* PE. **(D)** Representative Sanger sequencing of pgRNA-edited K052 clones showing sequence deletion targeting the *EZH2* PE. **(E)** Phase-contrast images of control (empty vector) or pgRNA-edited K562^Mut^ clones. **(F)** Representative gel showing the genomic PCR of control (empty vector) or pgRNA-edited K562^Mut^ clones targeting the *EZH2* PE. **(G)** Representative Sanger sequencing of pgRNA-edited K562^Mut^ clones showing sequence deletion targeting the *EZH2* PE.

### Genomic deletion of the *EZH2* PE corrects aberrant splicing and restores protein expression in SRSF2^Mut^ cancer cells

We next investigated the effect of deletion of the poison exon on splicing and protein expression. RT-PCR of extracted RNA from pgRNA-edited K052 cells showed loss of poison exon inclusion from *EZH2* mRNA as we expected (**Fig. 5A**). Western blotting of extracted protein also showed significant upregulation of EZH2 protein levels in pgRNA-edited K052 cells (**Fig. 5B**), supporting our hypothesis. We also performed RT-PCR and western blotting on pgRNA-edited K562^Mut^ cells and observed similar results for both splicing and protein expression (**Fig. 5C-D**). Taken together, pgRNA-mediated genomic excision of PE can reliably correct aberrant AS-NMD and restore protein expression.

**Figure 5.**
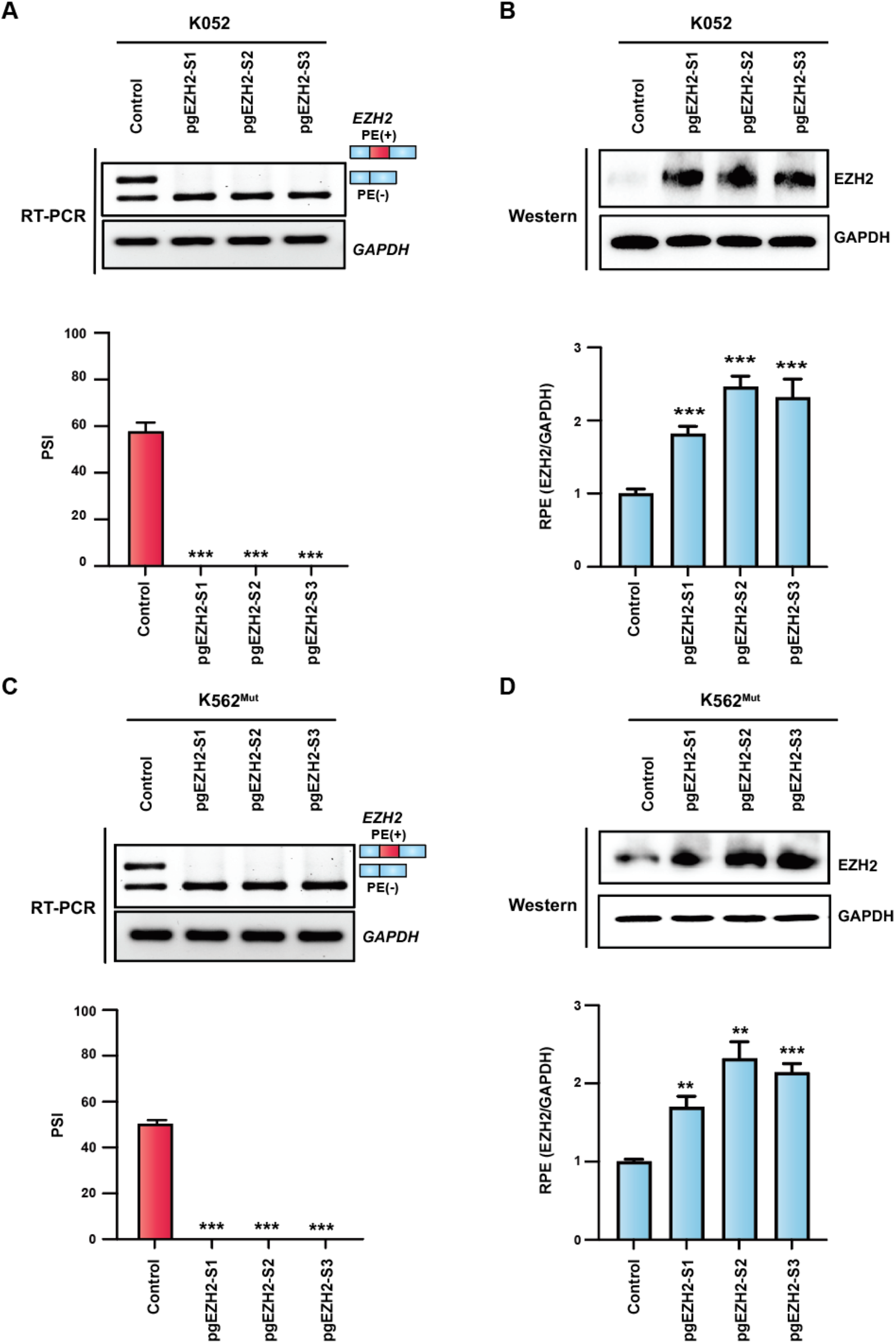
pgRNA-mediated genomic deletion of the *EZH2* PE corrects aberrant splicing and restores EZH2 protein expression in SRSF2-mutated model cancer cell lines. **(A)** Top, representative RT-PCR gels of control (empty vector) or pgRNA-edited K052 clones showing the splicing profiles of the *EZH2* PE. PE (+): PE inclusion; PE (-): PE exclusion. Bottom, quantification of the *EZH2* PE inclusion in mRNA as percent spliced-in (PSI). Bar graph (mean ± SD, n=3, ***p<0.001, t test). (**B**) Top, representative western blotting of control (empty vector) or pgRNA-edited K052 clones using antibodies against EZH2 and GAPDH. Bottom, quantification of the relative protein expression (RPE) of EZH2 normalized against GAPDH (EZH2/GAPDH). Bar graph (mean ± SD, n=3, ***p<0.001, t test). (**C**) Top, representative RT-PCR gels of control (empty vector) or pgRNA-edited K562^Mut^ clones showing the splicing profiles of the EZH2 PE. Bottom, quantification of the EZH2 PE inclusion in mRNA as PSI. Bar graph (mean ± SD, n=3, ***p<0.001, t test). (**D**) Top, representative western blotting of control (empty vector) or pgRNA-edited K562^Mut^ clones using antibodies against EZH2 and GAPDH. Bottom, quantification of the relative protein expression (RPE) of EZH2 normalized against GAPDH (EZH2/GAPDH). Bar graph (mean ± SD, n=3, **p<0.01, ***p<0.001, t test).

### *EZH2* poison exon deletion enhances histone methylation and rescues defective chromatin regulation in SRSF2^Mut^ cancer cells

We next seek to determine whether pgRNA-mediated restoration of EZH2 protein expression can rescue downstream molecular maladies. EZH2 has been implicated in chromatin regulation as part of the polycomb repressing complex 2 (PRC2). PRC2 functions in silencing gene expression through chromatin modification, playing a crucial role in stem cell differentiation and development (49). The functional core of PRC2 includes three nonenzymatic subunits, EED, SUZ11, and RBBP4, and one of the two enzymatic subunits (EZH2 or EZH1) (**Fig. 6A**) (49, 53). PRC2 catalyzes methylation of histone H3 at lysine 27 (H3K27me3), operating as a “writer” that labels chromatin for silencing. In a murine model, the Pro95 mutation in *Srsf2* was shown to promote aberrant splicing of *EZH2* and lead to MDS with impaired hematopoietic differentiation, peripheral cytopenias, and morphologic dysplasia (25). Furthermore, global H3K27 methylation was downregulated with EZH2 downregulation in SRSF2^Mut^ cells derived from a leukemia patient. It was also shown that restoring EZH2 protein expression using a cDNA partially rescues hematopoiesis in c-Kit+ cells isolated from the *Srsf2*-mutant murine model (25). We next measured total histone H3 and H3K27me3 protein levels in control and pgRNA-treated K052 cells. The results showed that the overall expression levels of histone H3 were similar in pgRNA-treated K052 and control K052. In contrast, H3K27me3 levels increased significantly in pgRNA-edited K052 compared to control K052 (**Fig. 6B**). To confirm that the increased H3K27me3 levels were solely due to increased EZH2 levels, we treated pgRNA-edited K052 cells with an EZH2 inhibitor (EZH2i) (**Fig. 6B**). As expected, treatment with the EZH2 inhibitor downregulated the H3K27me3 levels, confirming the EZH2-mediated H3K27 methylation (**Fig. 6B**).

**Figure 6.**
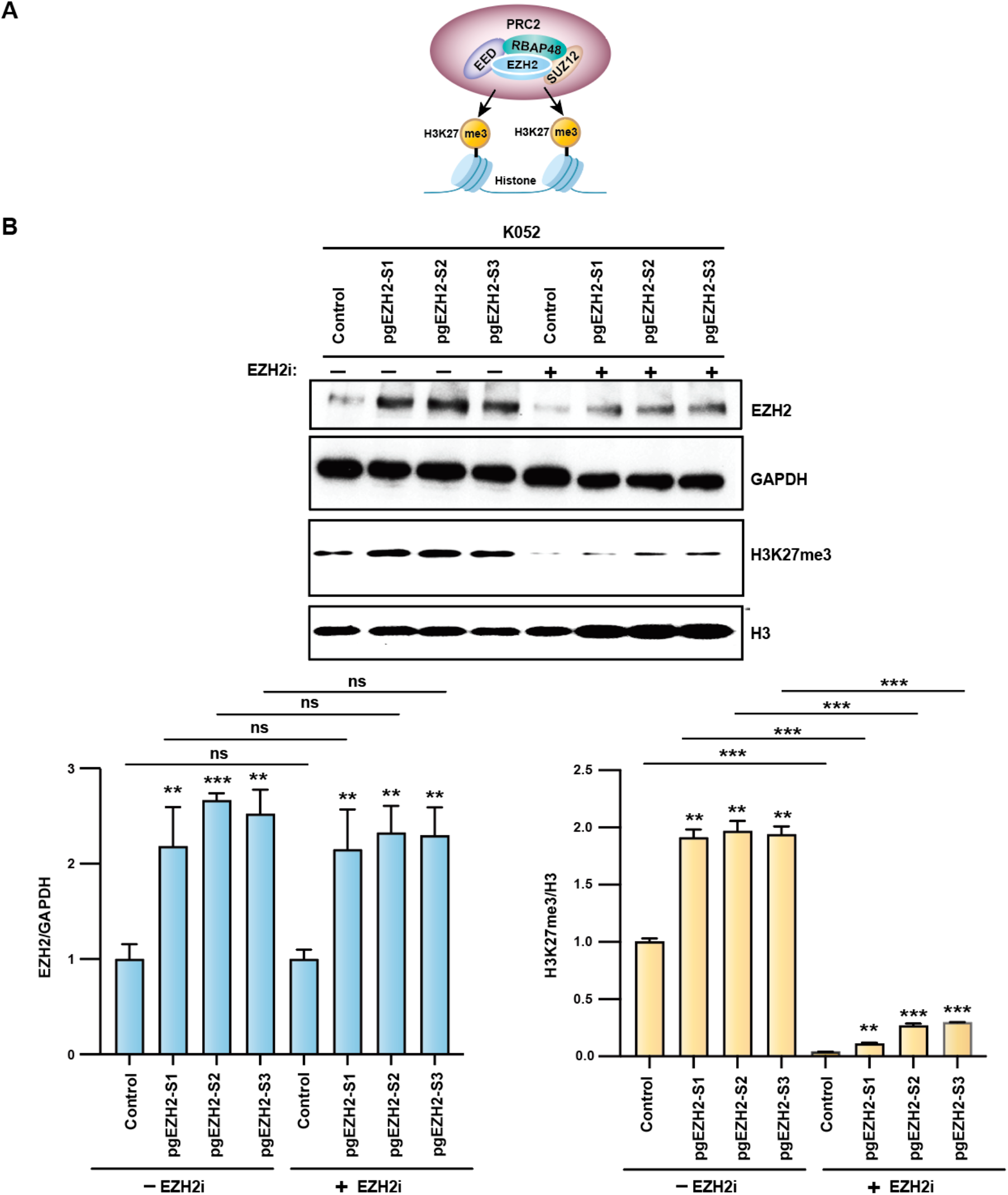
pgRNA-mediated genomic deletion of the *EZH2* PE rescues defective chromatin regulation in SRSF2-mutated model cancer cell line. **(A)** Diagram showing the core components of polycomb repressing complex (PRC2) and subsequent histone methylation at H3K27. **(B)** Top, representative western blotting of control (empty vector) or pgRNA-edited K052 clones treated without (-) or with (+) an EZH2 inhibitor (EZH2i) using the indicated antibodies. Bottom, quantification of the relative protein expression of EZH2 (EZH2/GAPDH) and H3K27me3 (H3K27me3/H3). Bar graph (mean ± SD, n=3, **p<0.01, ***p<0.001, ns: not significant, ANOVA).

### Comparative study between the proposed CRISPR strategy and antisense technology for modulating the oncogenic splicing of *EZH2*

We recently developed an antisense oligonucleotide candidate (ASO2) targeting a splicing regulatory cis-element in the *EZH2* PE, which can inhibit exon inclusion and produce a canonical, functionally active exon-skipped *EZH2* isoform in SRSF2^Mut^ cancer cells (38). To compare the relative efficacy of the proposed CRISPR and antisense approaches, we performed side-by-side experiments in K052 (**Fig. 7A**). We used only one pgRNA-edited K052 (pgEZH2-S1) in this experiment. Based on our prior analyses (38), we used 100 nm ASO2 with locked nucleic acid (LNA) modifications in the ribose sugars. RT-PCR of extracted RNA disclosed that ASO2 treatment promoted significant skipping of the *EZH2* PE compared to control K052 (**Fig. 7B**), but to a lesser extent than pgEZH2-S1 (**Fig. 7C**). Western blotting of extracted protein also showed higher efficacy for pgEZH2-S1-edited K052 compared to ASO2-treated K052 in restoring EZH2 protein expression, and subsequent upregulation of H3K27me3 levels, consistent with mRNA data (**Fig. 7D-E**). One critical factor that can potentially affect the efficacy of the antisense strategy is the shelf life. Since splicing modulation by ASO is regulated by binding to the target sequence in the RNA, a reduction of ASO concentration over time may result in lower efficiency in splicing modulation and restoring EZH2 protein expression. Therefore, we performed a time-course analysis over 10 days following ASO2 transfection in K052 (**Fig. 8A-B**). We observed a reduction in efficiency after 6 days for both RNA and protein levels (**Fig. 8A-B**). In contrast, a similar time-course analysis in pgEZH2-S1-edited K052 showed no considerable changes over 10 days (**Fig. 8C-D**). Taken together, the CRISPR strategy shows preferential advantages over the antisense approach for modulating oncogenic AS-NMD of *EZH2* in SRSF2^Mut^ cancer cells.

**Figure 7.**
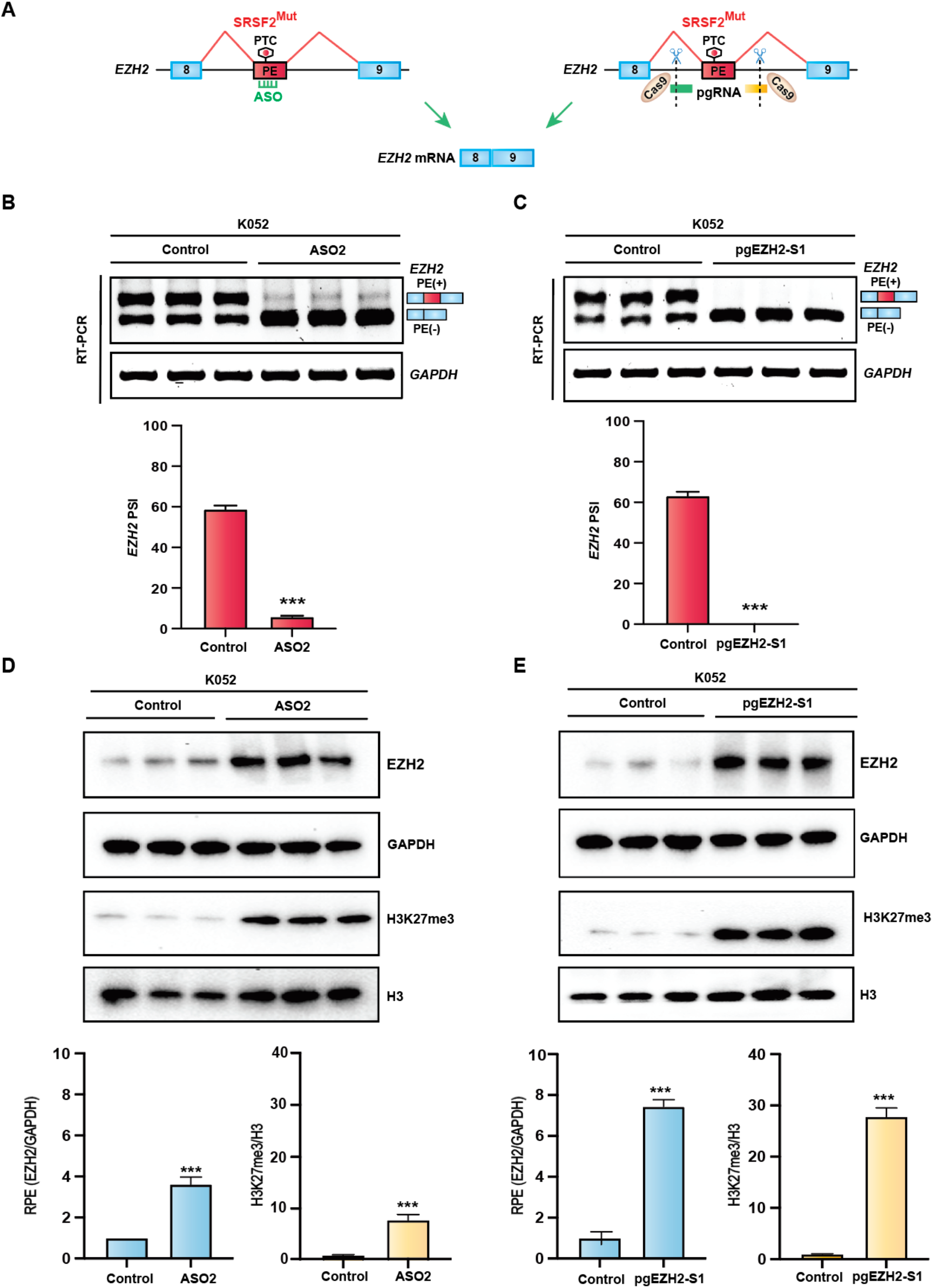
Evaluation of the relative efficacy between antisense pharmacology and CRISPR strategy. **(A)** Schematics showing antisense pharmacology and CRISPR strategy to modulate oncogenic PE inclusion of *EZH2* in SRSF2^Mut^ cancer cell line. **(B)** Top, representative RT-PCR gels of K052 cells showing the splicing profiles of the *EZH2* PE treated without (control) or with an antisense oligonucleotide candidate (ASO2). PE (+): PE inclusion; PE (-): PE exclusion. Bottom, quantification of the *EZH2* PE inclusion in mRNA as percent spliced-in (PSI). Bar graph (mean ± SD, n=3, ***p<0.001, t test). **(C)** Top, representative RT-PCR gels of control or pgRNA-edited K052 clones (pgEZH1-S1) showing the splicing profiles of the *EZH2* PE. Bottom, quantification of the *EZH2* PE inclusion in mRNA as PSI. Bar graph (mean ± SD, n=3, ***p<0.001, t test). **(D)** Top, representative western blotting for samples in panel (**B**) using the indicated antibodies. Bottom, quantification of the relative protein expression (RPE) of EZH2 (EZH2/GAPDH) and H3K27me3 (H3K27me3/H3). Bar graph (mean ± SD, n=3, ***p<0.001, t test). (**E**) Top, representative western blotting for samples in panel (**C**) using the indicated antibodies. Bottom, quantification of the relative protein expression (RPE) of EZH2 (EZH2/GAPDH) and H3K27me3 (H3K27me3/H3). Bar graph (mean ± SD, n=3, ***p<0.001, t test).

**Figure 8.**
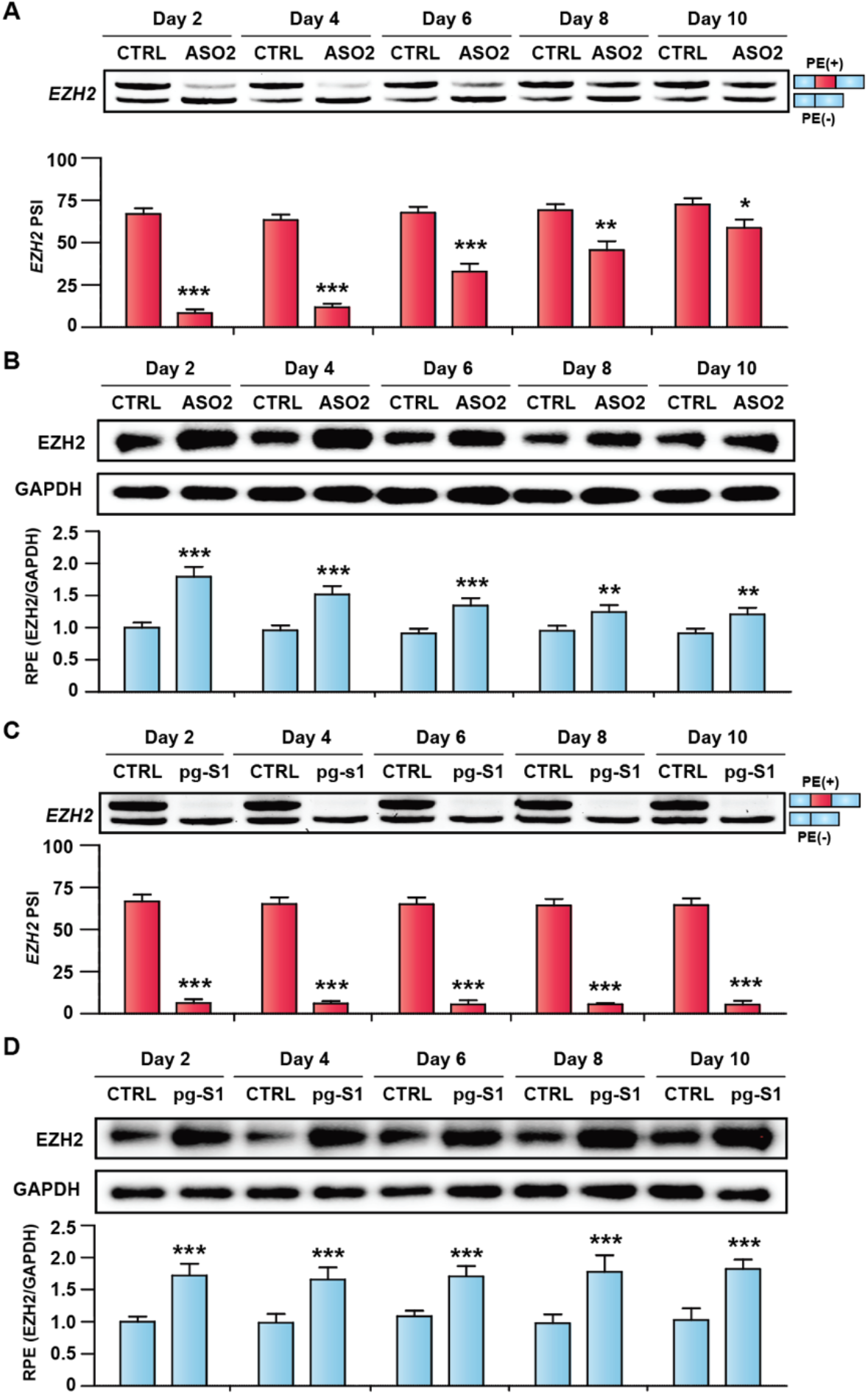
Time course profile of *EZH2* splicing and expression in SRSF2^Mut^ cancer cells treated with antisense oligonucleotide or pgRNA. **(A)** Top, representative RT-PCR gel of K052 cells showing a temporal profile of *EZH2* PE splicing treated without (CTRL) or with an antisense oligonucleotide candidate (ASO2). PE (+): PE inclusion; PE (-): PE exclusion. Bottom, quantification of the *EZH2* PE inclusion in mRNA as percent spliced-in (PSI). Bar graph (mean ± SD, n=3, *p<0.05, **p<0.01, ***p<0.001, t test). **(B)** Top, representative western blotting for samples in panel (**A**) using the indicated antibodies. Bottom, quantification of the relative protein expression (RPE) of EZH2 (EZH2/GAPDH). Bar graph (mean ± SD, n=3, **p<0.01, ***p<0.001, t test). **(C)** Top, representative RT-PCR gel of control (CTRL) or pgEZH2-S1-edited K052 clones (pg-S1) showing a temporal profile of *EZH2* PE splicing. Bottom, quantification of the *EZH2* PE inclusion in mRNA as PSI. Bar graph (mean ± SD, n=3, ***p<0.001, t test). (**D**) Top, representative western blotting for samples in panel (**C**) using the indicated antibodies. Bottom, quantification of the relative protein expression (RPE) of EZH2 (EZH2/GAPDH). Bar graph (mean ± SD, n=3, ***p<0.001, t test).

## Discussion

Since the milestone discovery of recurrent mutations in splicing factors in hematologic malignancies (23), the field has made extensive progress in unraveling molecular maladies by identifying aberrantly spliced targets, defective pathways, and mechanistic features. Although initial speculation suggested that mutations in splicing factors might affect the splicing of common downstream targets and pathways, genome-wide RNA sequencing results ruled out this hypothesis and demonstrated that distinct splicing factor mutations target distinct targets and pathways. This appeared to be a critical roadblock to developing a widely adopted strategy for treating all splicing-factor-mutant tumors. Despite these challenges, several therapeutic approaches have been exploited to date to treat cancer with splicing defects. These include targeting chemical compounds that bind core spliceosome components, splicing regulatory kinases (such as splicing factor kinases, SRPKs, and CLKs), protein arginine methyltransferases (PRMTs), and RNA-binding proteins (4, 5, 31, 54). However, those efforts were mostly unsuccessful due to off-target effects or an unclear mode of action. These results suggested that gene-specific targeted splicing modulation of aberrantly regulated genes, or selective inhibition of the function of specific mutant splicing factors, could be more effective strategies to avoid off-target effects. Although new strategies are emerging, such as synthetic introns (55) and oligonucleotide pharmacology (28) to selectively target splicing-factor-mutant cancer cells, therapeutic success remains elusive due to various technological shortcomings, including delivery, dose, stability, and throughput.

To push the boundaries and challenge existing paradigms in the field, we present a proof-of-principle demonstrating that CRISPR/Cas9 can restore the expression of crucial proteins degraded by PE-mediated AS-NMD and linked to tumorigenesis in SRSF2-Mutant cancer, using an efficient, gene-specific targeting strategy. We targeted the PE-mediated AS-NMD event in *EZH2*. It was reported that EZH2 is a crucial epigenetic regulator, and frequently dysregulated in MDS, AML, and other cancers (25–27, 49–51, 56, 57). Our strategy uses a single vector encoding pgRNA and Cas9 to stably delete the *EZH2* PE from the genome of SRSF2^Mut^ cancer cells. Our approach efficiently corrects aberrant PE inclusion, escapes AS-NMD, restores EZH2 protein expression, and rescues dysregulated chromatin regulation. Although CRISPR/Cas9 is typically used to knock out specific genes, our approach works in the opposite direction, restoring gene expression by modulating splicing at an alternative cassette exon. It is important to note that CRISPR/Cas9-mediated genome editing often results in the insertion or deletion of nucleotides in the host genome, generating PTCs in mRNA, and subsequently downregulates gene expression through NMD. However, our approach is likely to avoid this problem as our pgRNA is designed to target the intronic sequences flanking the alternative cassette exon. Therefore, even if nucleotides are inserted or deleted in the host genome during the editing process, they will be excluded during intron splicing without disrupting the mRNA frame.

We recently developed antisense oligonucleotides (ASO) targeting an exonic splicing enhancer (ESE) element of the *EZH2* PE, which can prevent aberrant PE inclusion and AS-NMD, and successfully restore EZH2 protein expression. However, the ASO can’t promote complete splice-switching to generate an exon-skipped productive EZH2 isoform, which is potentially affected due to suboptimal delivery or dose of ASO and subsequent binding to the ESE element in the target exon. Furthermore, the stability and shelf life of ASO are additional factors that can affect splice-switching efficacy after administration. Compared to ASO, CRISPR/Cas9-mediated splicing modulation showed more efficient splice-switching as it permanently deleted the PE from the genome. However, this approach is applicable only to alternative cassette exons. For other AS-NMD events arising from skipping of an alternative exon, inefficient intron splicing, or alternative splice-site selection, we need to use an alternative CRISPR/Cas9 approach, such as mutating or deleting critical cis-elements that regulate splicing. Our future studies will focus on developing such novel strategies.

Mutations in splicing factors promote genome-wide splicing alterations; however, only several altered genes are linked to tumorigenesis (cancer drivers). For example, in SRSF2-mutated cancer, aberrant AS-NMD in *EZH2, INTS3, and CLK3* were experimentally validated with a proven link to tumorigenesis. Therefore, correcting AS-NMD in a single gene is expected to partially rescue defective hematopoiesis and confer anticancer effects, which is a limitation for cancer therapy compared with monogenic genetic diseases. To address this shortcoming, we hope to leverage advances in CRISPR technology to develop a strategy that simultaneously corrects AS-NMD in multiple genes from a single vector, achieving more beneficial outcomes through combinatorial effects. Although the focus of this study was to develop a therapeutic proof-of-concept targeting a representative oncogenic AS-NMD event promoted by poison exon, our future research will develop a novel CRISPR strategy for combinatorial therapy targeting multiple genes and will test it in preclinical animal models *in vivo,* with the hope of advancing to clinical development.

## Acknowledgments

We are grateful to Omar Abdel-Wahab (Memorial Sloan Kettering Cancer Center) for sharing human leukemia cell lines. A few diagrams were created with BioRender.

## Author contributions

M.A.R. conceived the project. M.A.R. supervised the study. M.A.R., M.H., P.N., and M.R.I. designed experiments. P.N. and M.R.I. performed all the experiments with the help of N.A.R., S.A.H., M.M.H., S.H., and R.A.; P.N., M.R.I., and M.A.R. wrote the primary manuscript with inputs from all authors. M.A.R. and M.H. finalized the manuscript. All authors approved the final version of the manuscript.

## Conflict of interest

M.A.R., M.R.I., and P.N. are inventors on a patent application not related to this study. Other authors declare no conflict of interest.

## Funding

This work was supported by the National Institute of General Medical Sciences (NIGMS) of the National Institutes of Health (R35GM154991 to M.A.R.) and the Winthrop P. Rockefeller Cancer Institute.

